# Novel insights into the structure and transport mechanisms of TAPT1

**DOI:** 10.1101/2020.05.18.099887

**Authors:** Md Sorwer Alam Parvez, Mohammad Mahfujur Rahman, Md Niaz Morshed, Dolilur Rahman, Saeed Anwar, Paul J. Coucke, Mohammad Jakir Hosen

**Affiliations:** Department of Genetic Engineering & Biotechnology, Shahjalal University of Science & Technology, Sylhet 3114, Bangladesh; Department of Medical Genetics, Faculty of Medicine and Dentistry, University of Alberta, 8440 112 St. NW, Edmonton, AB T6G 2R7, Canada; Center for Medical Genetics, Ghent University Hospital, 9000 Ghent, Belgium

**Keywords:** TAPT1, topology modeling, substrate-binding site, homology modeling, molecular docking

## Abstract

Transmembrane anterior-posterior transformation protein 1 (TAPT1), encoded by the *TAPT1* gene expressed in the basal ciliary body, plays a crucial role in cilia formation as well as axial skeletal patterning. Mutations in this gene have been reported to cause several ciliopathies and osteo-related diseases. Unfortunately, the cellular and molecular pathogenic mechanisms are still unclear also due to the lack of X-ray crystallographic structure and further characterization of TAPT1 protein. In this study, we attempted to characterize this protein by *in silico* techniques. A 3D structure of TAPT1 was generated by the *ab initio* method, which was further used for the analysis of the substrate-binding site, to determine pore size and for the prediction of the possible substrate(s). Validation by using different software packages revealed a reliable 3D model of TAPT1. Topology modeling revealed that TAPT1 has eight transmembrane helices with a total number of 27 helices in secondary structure. The amino acid residues H235, R323, K443, N446, S447, L450, K453, S454, Y457, K511, N513, D533, K535, D536, and T538 were found to form the pore surface as well as involved in the binding interaction with the substrate(s). This study predicted flavonoids as the possible substrate for TAPT1, which could further be confirmed by ingenuity pathway analysis. Moreover, our analysis indicated that TAPT1 might localize in the mitochondrial membrane in addition to the ciliary basal body. Our study gives novel insights for TAPT1 structure and its function.

## 1. Introduction

The evolutionary conserved human trans-membrane anterior-posterior transformation protein 1 (TAPT1) is encoded by the TAPT1 gene, located at Ch4 p15.32. In eukaryotes, TAPT1 plays a role in axial skeletal patterning during early embryonic development [1, 2]. Mutations in TAPT1 causes a lethal congenital syndrome, by which clinical features overlap between skeletal dysplasia and ciliopathies [3]. The cellular and molecular mechanisms causing the syndrome are still unclear. Although expression of the TAPT1 is noticed at most body sites, localization is most common at the centrosome and basal ciliary body [4]. Defective TAPT1 mislocalized in the cytoplasm disrupts the morphology of the Golgi apparatus and intracellular protein trafficking, giving rise to abnormal cilia formation [3, 5].

In mice, Tapt1 is predicted to comprise of several transmembrane domains, and part of the gene is orthologous to alternatively spliced human transcript encoding the cytomegalovirus gH receptor. Besides, Howell et al. also speculated that Tapt1 acts as a downstream effector of Hoxc8 that may transduce extracellular information required for axial skeletal patterning during embryonic development [6]. In zebrafish models, knockdown of tapt1b revealed delayed ossification and abnormal craniofacial skeleton development, linked to abnormal cranial neural crest cell differentiation [3].

To date, several studies endeavored to characterize TAPT1. However, the absence of a high-resolution X-ray crystallographic structure, the unknown substrate(s), and undefined molecular signaling pathway challenged the characterization of the protein. A comprehensive in silico analysis, including 3D homology modeling and other computational approaches, could be an alternative approach to gain insights into the proteins’ structure, potential binding sites, and the possible substrate(s). Such studies would help gain envisage the mechanisms of substrate interaction with the protein, as well as to disclose involved molecular signaling pathways [7–9]. The current study aimed to characterize TAPT1, both structurally and functionally, using computational approaches, which may give novel insights into the function of this protein and understand the underlying pathogenic mechanisms of the diseases associated with it.

## 2. Materials and Methods

### 2.1 Prediction of secondary structure and topology model of TAPT1

The primary sequence of human TAPT1 protein (accession no: AAH66899.1) was retrieved from the GenBank database of the National Center for Biotechnology Information (NCBI), which has a sequence length of 567 amino acids [10]. The protein secondary structure was analyzed using PSIPRED secondary structure prediction server, and the topology was predicted by the MEMSAT-SVM of the PSIPRED workbench online server [11–13]. PDBsum was also used to predict the secondary structure from the generated 3D model [14].

### 2.2 Generation and validation of TAPT1 3D homology model

The Basic Local Alignment Search Tool (BLAST) program of NCBI was used against the Protein Data Bank (PDB) to find the template for the homology modeling [15]. Unfortunately, no significant hit was found. Henceforth, the *ab initio* method was used to generate the 3D model of TAPT1. The Robetta server was used to model TAPT1 by the *ab initio* method [16]. Further, the generated 3D model was validated using RAMPAGE, PROCHECK, and ERRAT [17–19]. For the visualization of the model, a user-sponsored molecular visualization system, PyMOL was used [20].

### 2.3 Determination of pore diameter and pore-lining residues

We analyzed the transporter channel using the PoreWalker server, a fully automated method designed to detect and characterize transmembrane protein channels from their 3D structure [21]. Further, the PoreLogo server was used to visualize the sequence and conservation of pore-lining residues in transmembrane protein structures [22].

### 2.4 Subcellular localization and transporter substrate specificity prediction

The subcellular localization of this transporter was predicted by LocTree3 server which predicts the subcellular localization for all proteins in all domains of life where Water-soluble globular and transmembrane proteins are predicted in one of 18 classes in Eukaryota (e.g., chloroplast, chloroplast membrane, cytosol, ER, Golgi, ER membrane, Golgi membrane, extracellular, mitochondria, mitochondria membrane, nucleus, nucleus membrane, peroxisome, peroxisome membrane, plasma membrane, plastid, vacuole, and vacuole membrane) [23]. Further, the TrSSP server was used to predict the types of substrates specific for this transporter. This server predicts the substrate specificity based on evolutionary information and the AA index [24].

### 2.5 Protein-protein interaction analysis and prediction of the functions

GeneMANIA was used for the protein-protein interaction analysis. These tools also predict the functions by analyzing a large number of functional association data, including protein and genetic interactions, pathways, co-expression, co-localization, and protein domain similarity [25].

### 2.6 Molecular docking analysis for substrate prediction

Molecular docking analysis was carried out by AutoDock vina against predicted specific types of metabolites or molecules reported in the Human Metabolome Database (HMDB) [26, 27]. In our previously mentioned method, TrSSP predicted that TAPT1 involved in the transportation of charged molecules or metabolites. For which, only charged molecules or metabolites of HMDB was used as target ligands for molecular docking analysis. During this docking analysis, the grid box size was set to 75, 100, and 75 respectively for X, Y, and Z-axis, where the center was set to 35.254, 3.105 and 22.549 respectively for X, Y, and Z. Further, the binding site was analyzed in the PyMOL, and additionally, CASTp was used to cross-check the biding sites [28].

### 2.7 Ingenuity pathway analysis for possible pathway prediction

The top predicted substrate was then used for the ingenuity pathway analysis against TAPT1 by Ingenuity Pathway Analysis (IPA) [29]. IPA is an advanced bioinformatics tool that can generate the shortest possible pathway for the interesting data sets through ingenuity knowledge-based or text mining analysis. The shortest possible pathways were generated using the path explorer features of IPA, where the predicted substrate was selected as initial molecules. A hypothetical pathway was illustrated with Biorender.com concluding all the findings in this study.

## 3. Results

### 3.1 Generation and validation of human TAPT1 *ab initio* model

Mining of Protein Data Bank (PDB) revealed no high-resolution X-ray crystallographic structure and the homolog of TAPT1 is available. Thus, Robetta online server was used to generate a 3D homology model of TAPT1. Robetta generated five primary models, which were further validated using several structure assessment methods, including PROCHECK, ERRAT, and Ramachandran plot. Among the five, the best model (Fig. 1) was then selected based on the highest score of various parameters obtained from assessment methods. While the Ramachandran plot (generated by RAMPAGE) showed 99.5% residues of the best model are in the favored regions, PROCHECK showed 93.3% residues of this model were in most favored regions. ERRAT (which analyzes statistics of non-bonded interactions between different atom types based on characteristic atomic interaction) also showed an overall quality factor of the best-selected model is 97.967% (Fig. 2). Thus, the constructed best-selected model was highly reliable for further docking study.

**Fig 1:**
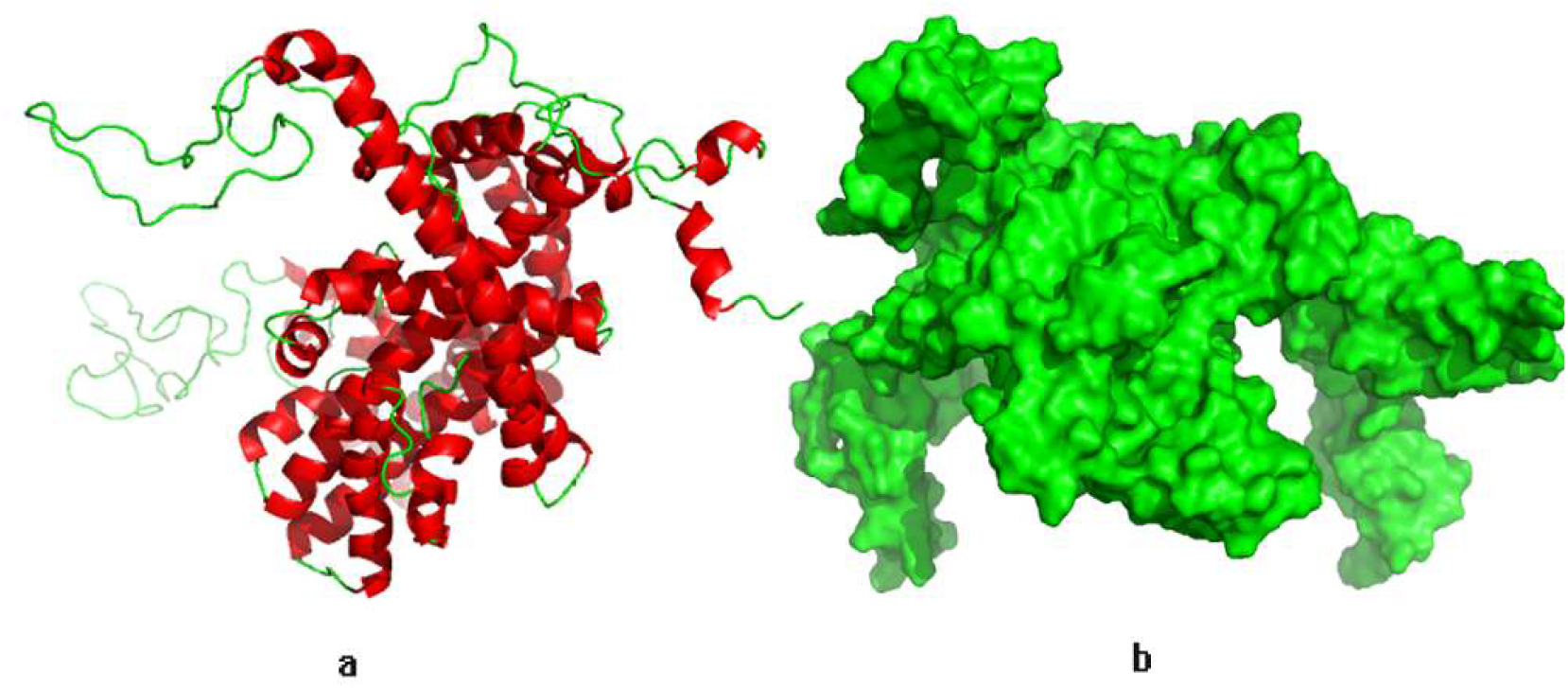
3D model of TAPT1. (a) The ribbon model where red color represents the helices and green represents the loop. (b) Surface model of TAPT1

**Fig 2:**
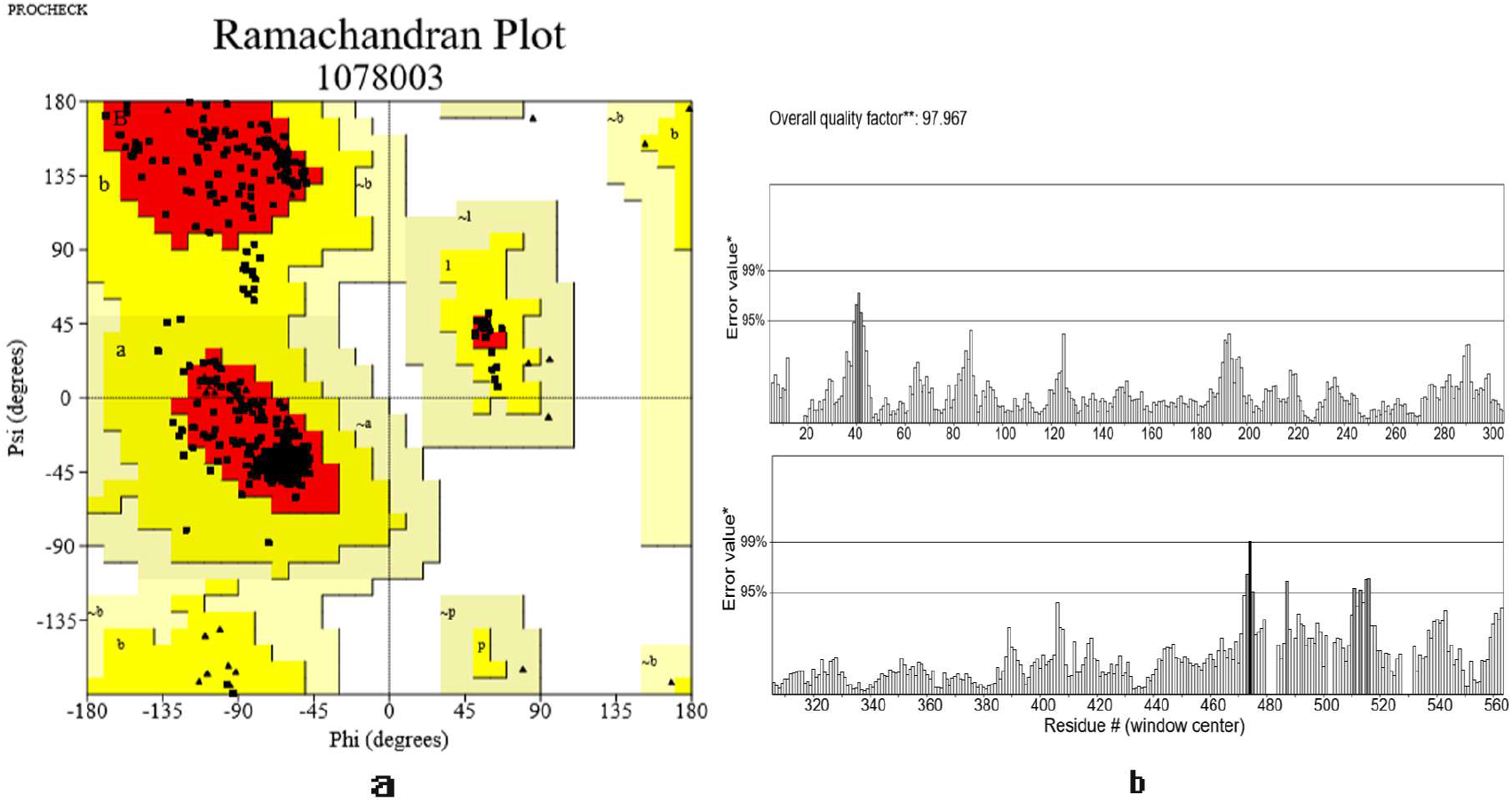
3D model validation assessment results where Ramachandran plot was shown in (a), and error value was shown in (b)

### 3.2 Secondary structure and topology model of TAPT1

MEMSAT-SVM mediated topology modeling predicted that TAPT1 comprised eight transmembrane helices, and both the N-terminal and C-terminal region located in the extracellular region (Fig. 3). The secondary structure of TAPT1 was predicted by the PDBsum, which revealed the presence of 27 helices, 32 beta turns, and a disulfide bond (Fig. 4).

**Fig 3:**
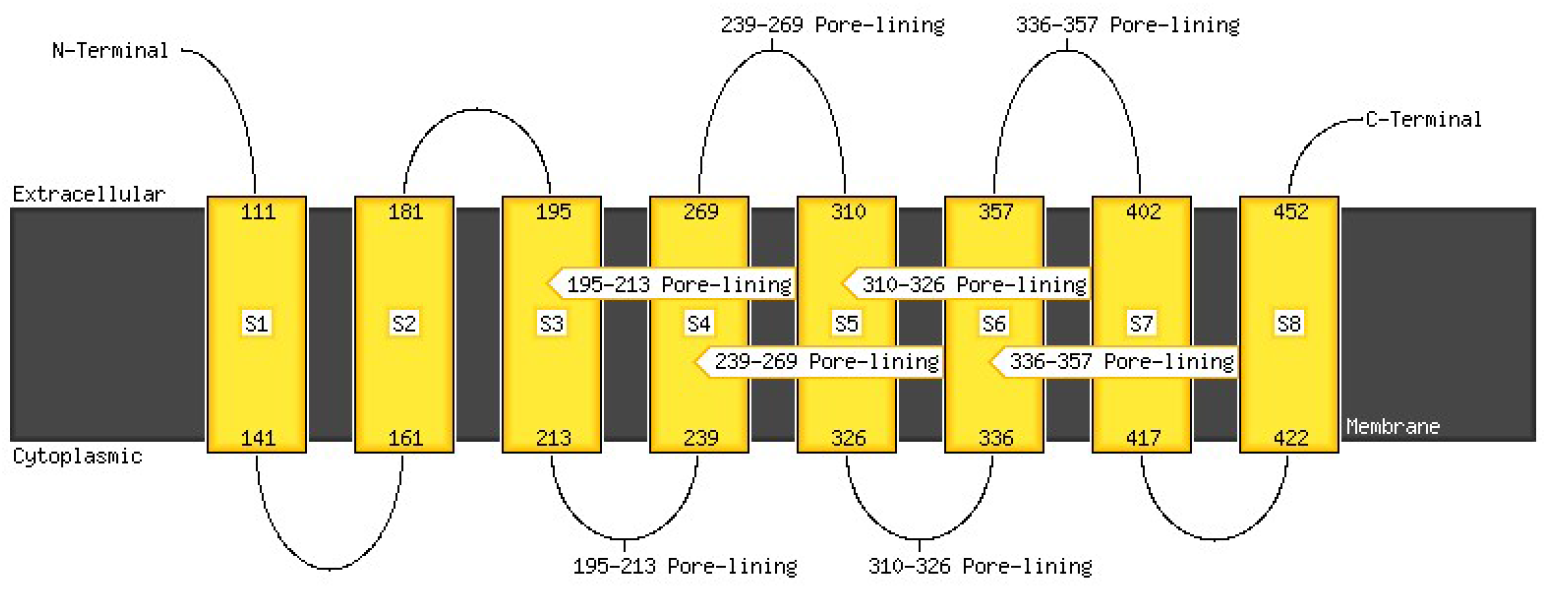
Topology of the TAPT1 protein

**Fig 4:**
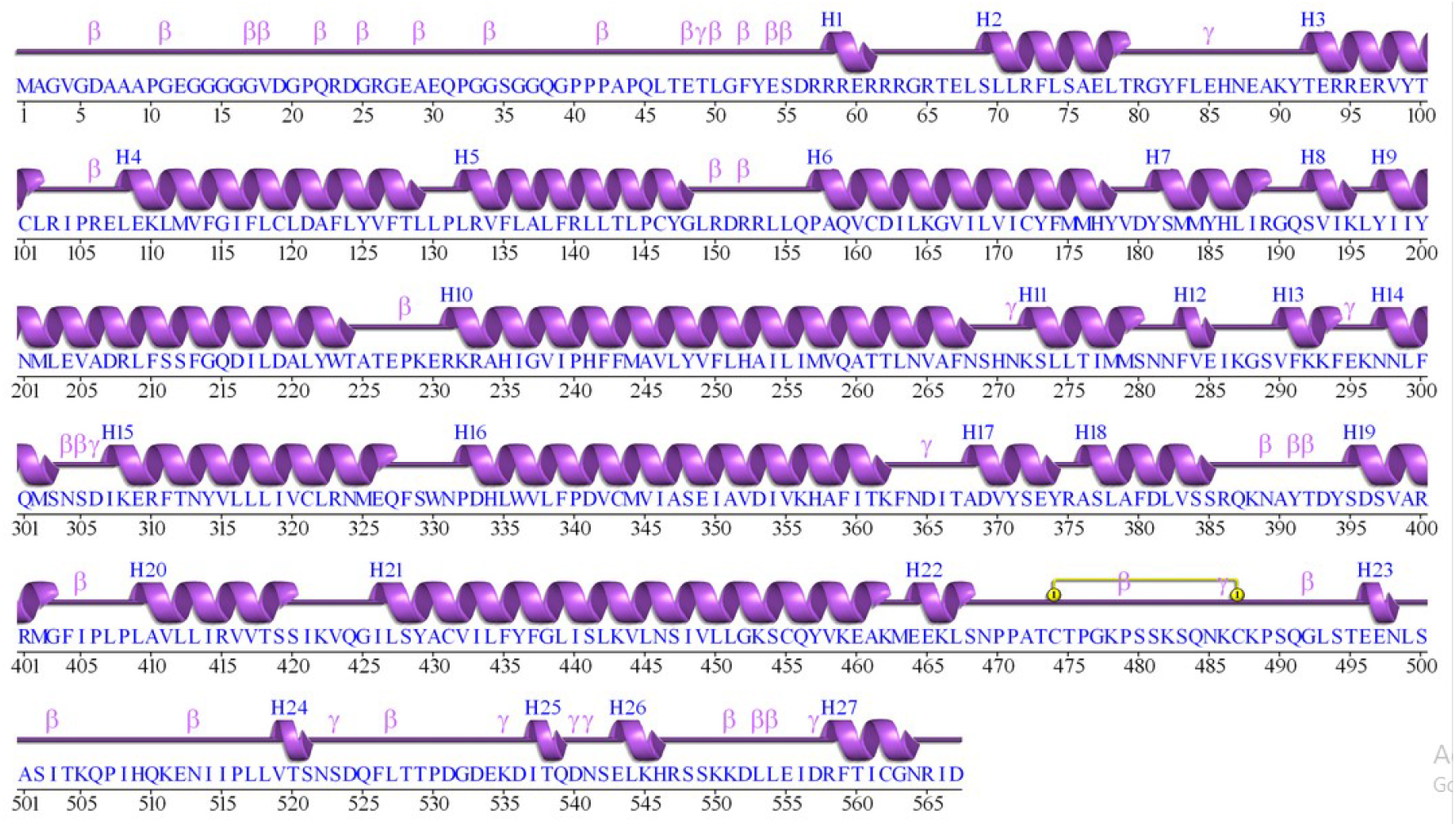
Secondary structure prediction from the 3D model where H represents the helices, β represents beta-turn, γ represents the gamma turn, and the yellow line represents the disulfide bond.

### 3.3 Insights of the TAPT1 transport channel pore

This study revealed that 105 amino acids were involved in the formation of transport pore of the TAPT1, where amino acid residues H235, R323, K443, N446, S447, L450, K453, S454, Y457, K511, N513, D533, K535, D536, and T538 were present in the pore surface (Fig 5). Moreover, the pore was found in a diamond shape.

**Fig 5:**
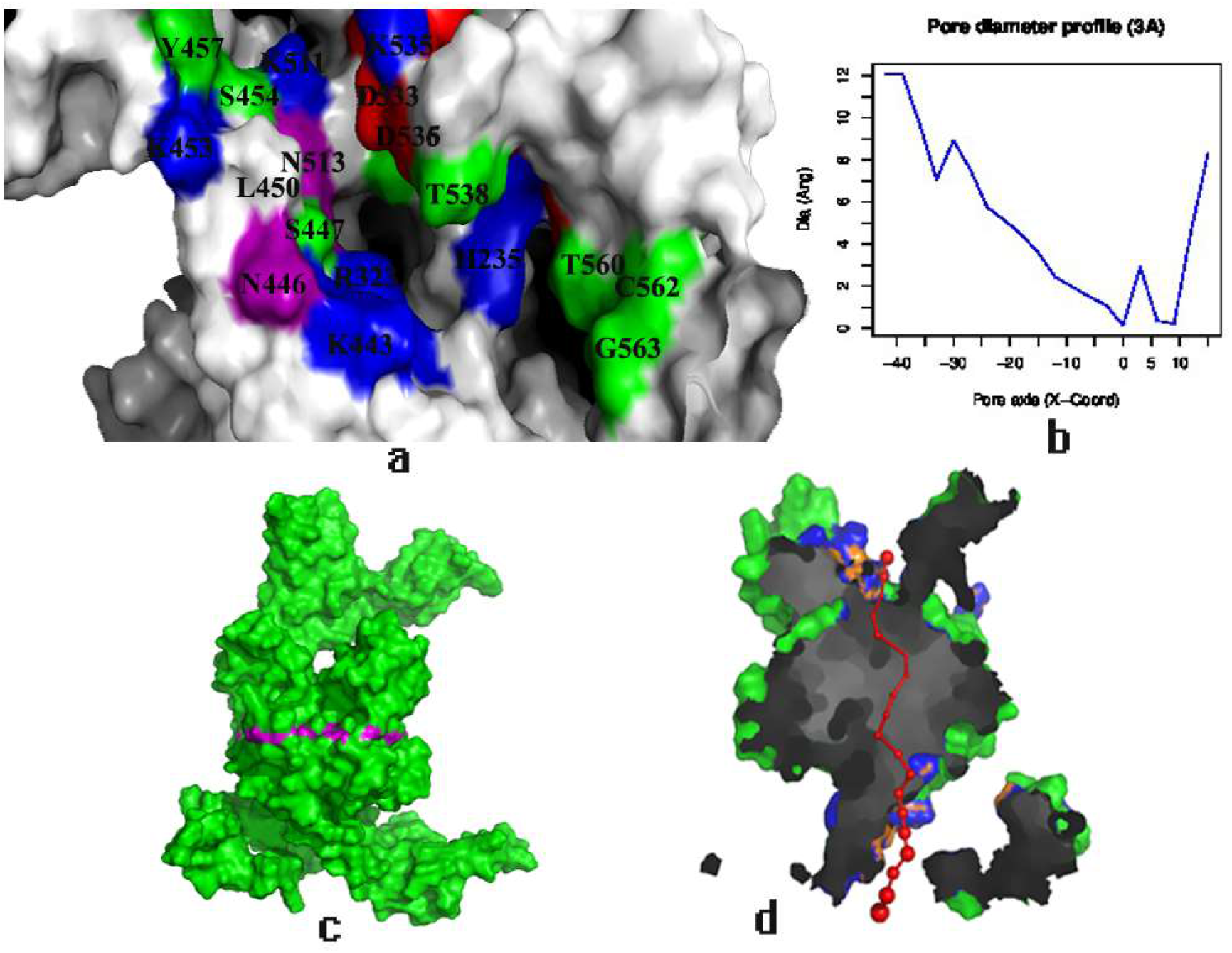
Protein-protein interactions of TAPT1. Here, pink lines represent physical interactions. Violet lines represent the co-expression, and gold lines represent the predicted interactions.

### 3.4 Interactome analysis

Interactomics revealed that TAPT1 has interaction with 20 other genes with a total number of 268 links. This gene has physical interactions with MIEF1, MYO10, SQRDL, P4HB, NDUFA13, DERL1, TXNDC15, SUCO, CAV1, RIC3, UBC, and PCNT while co-expressed with DDX6, NDUFA13, NAPA, SUCO, UBC, LYRM5, PTAR1, MED17and RBL2 (Fig 6). Moreover, this interactomics predicted that TAPT1 might involve in the regulation intrinsic apoptotic signaling pathway, cellular response to oxygen level, intracellular transportation of virus and viral life cycle, and multi-organism transportation.

**Fig 6:**
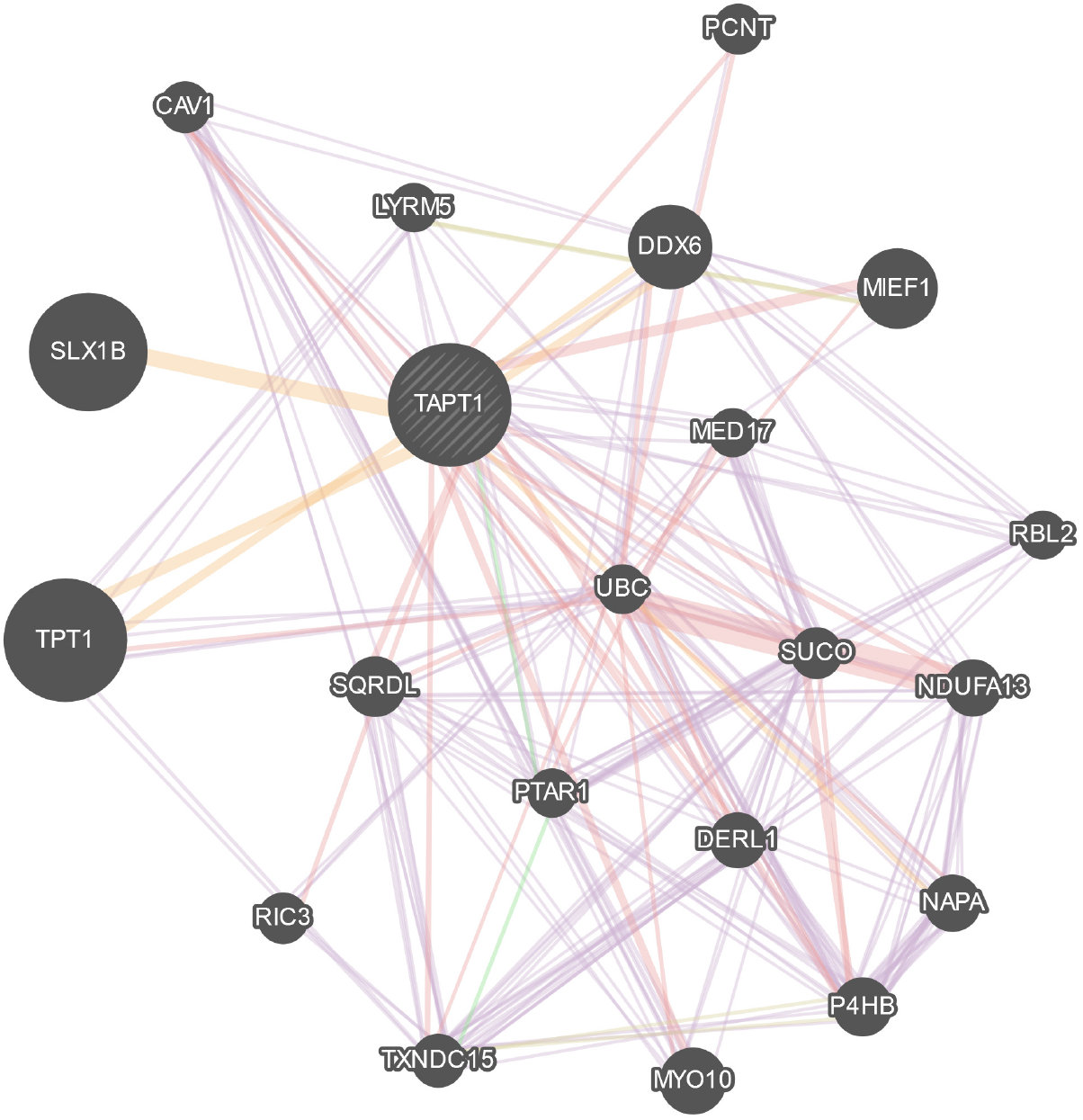
Pore of the TAPT1 where (a) shows the pore surface with involved residues (b) pore diameter profile in 3 Armstrong (c) pore line from the surface side and (d) pore line from top to bottom

### 3.5 Subcellular localization and substrate affinity of TAPT1

Prediction of subcellular localization via LocTree3 revealed that TAPT1 possibly localized in the mitochondrial membrane, and the expected accuracy was 84%. Besides, substrate specificity analysis stated that TAPT1 would involve in the transportation of charged (both positive and negative) metabolites.

### 3.6 Prediction of TAPT1 substrate by molecular docking and virtual screening

We screened all the charged metabolites reported in the Human Metabolome DataBase (HMDB) as target ligands of the TAPT1 model. Molecular docking analysis revealed that flavonoid glycosides (HMDB0124740) have the highest affinity with the lowest binding energy (Table 1). Surprisingly, all the top 10 compounds with the lowest binding energy were found in this subclass. Moreover, the binding site analysis revealed that amino acid A220, W223, T224, I239, R323, K443, S447, L450, N513, and P516 were involved in the interaction with the substrate (Fig 7). The same binding site was also predicted by the CASTp (Supplementary Fig 1).

**Table 1:**
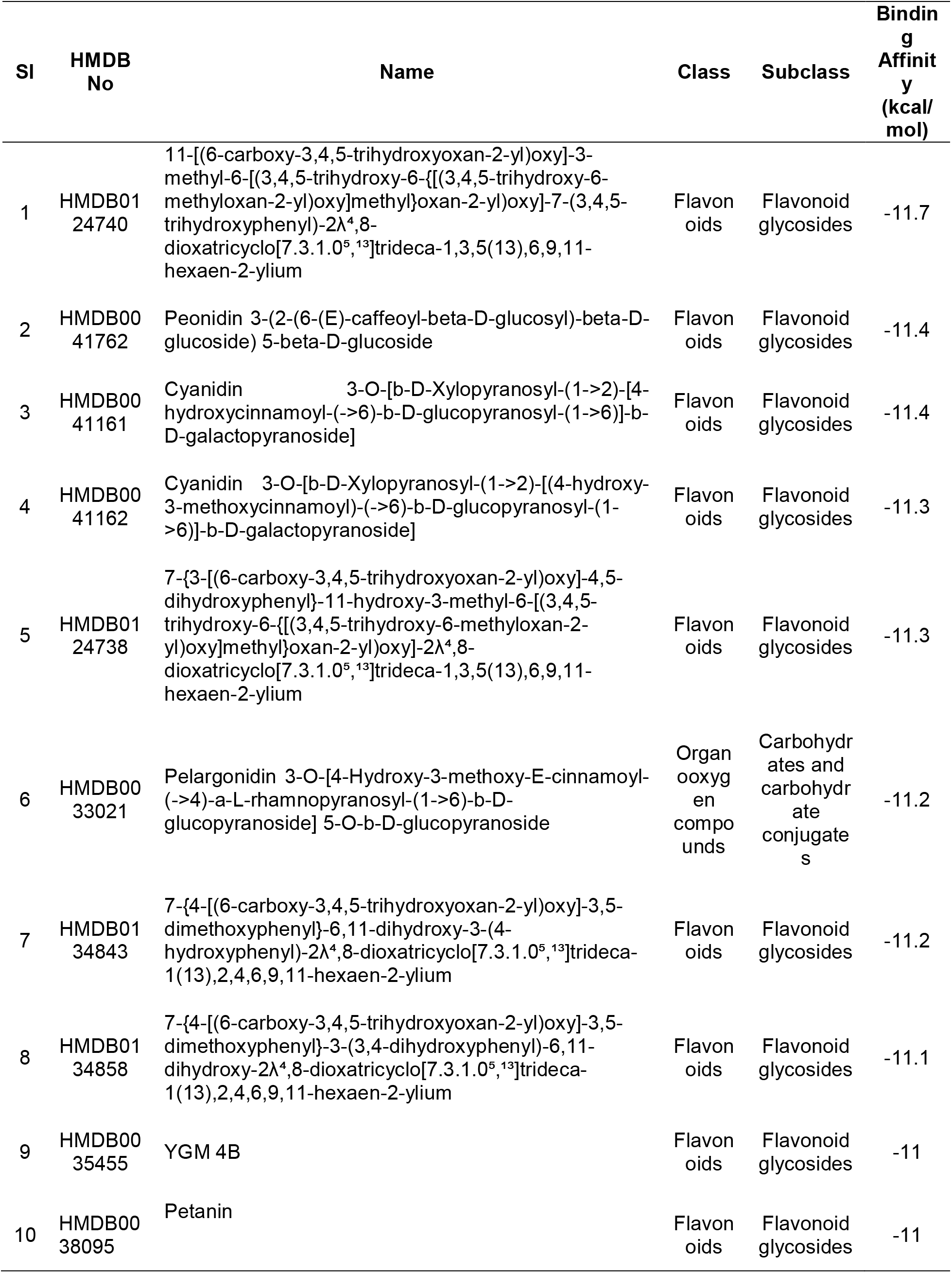
Top 10 compounds based on the highest binding energy against TAPT1

**Fig 7:**
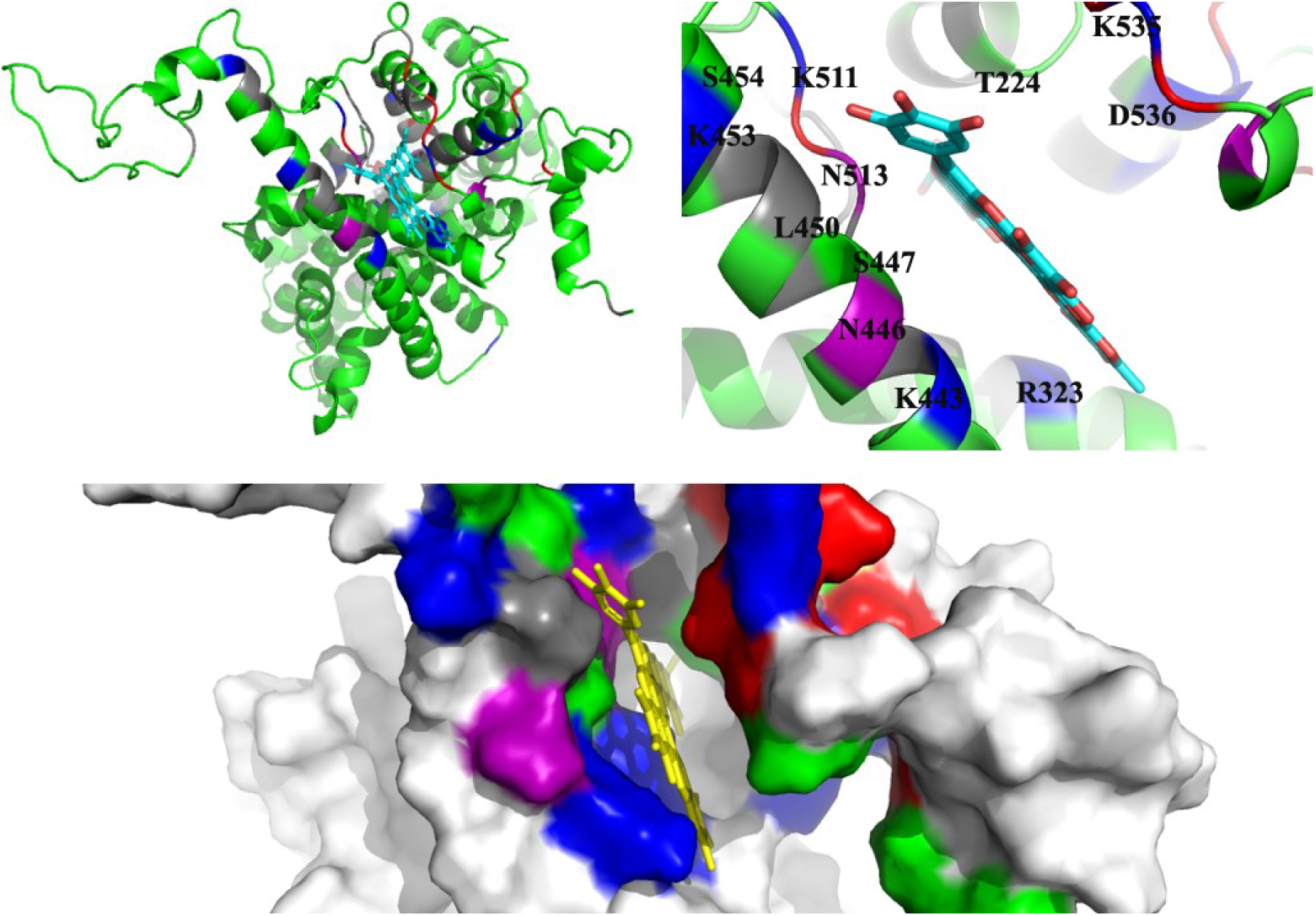
Docked figures of TAPT1 with its substrates. Here, different colors were used to represent the different pore-lining residues where blue color was used for Lys and His residues and red for Asp and Glu residues. Additionally, green used for Gly, Cys, Ser, Thr, and Tyr residues and gray for the remaining residues.

### 3.7 Involvement of TAPT1 in different signaling pathways

Ingenuity pathway analysis was performed to understand the relationship between flavonoid and TAPT1, which revealed that flavonoid induced the expression of several genes, including ERK1/2, AHR, HMOX1, CYP1A1/2, and estrogen receptor, which in turn induced CCND1 and CAV1. CCND1 increases the expression of TAPT1, where CAV1 binds to TAPT1 to activate it (Fig 8).

**Fig 8:**
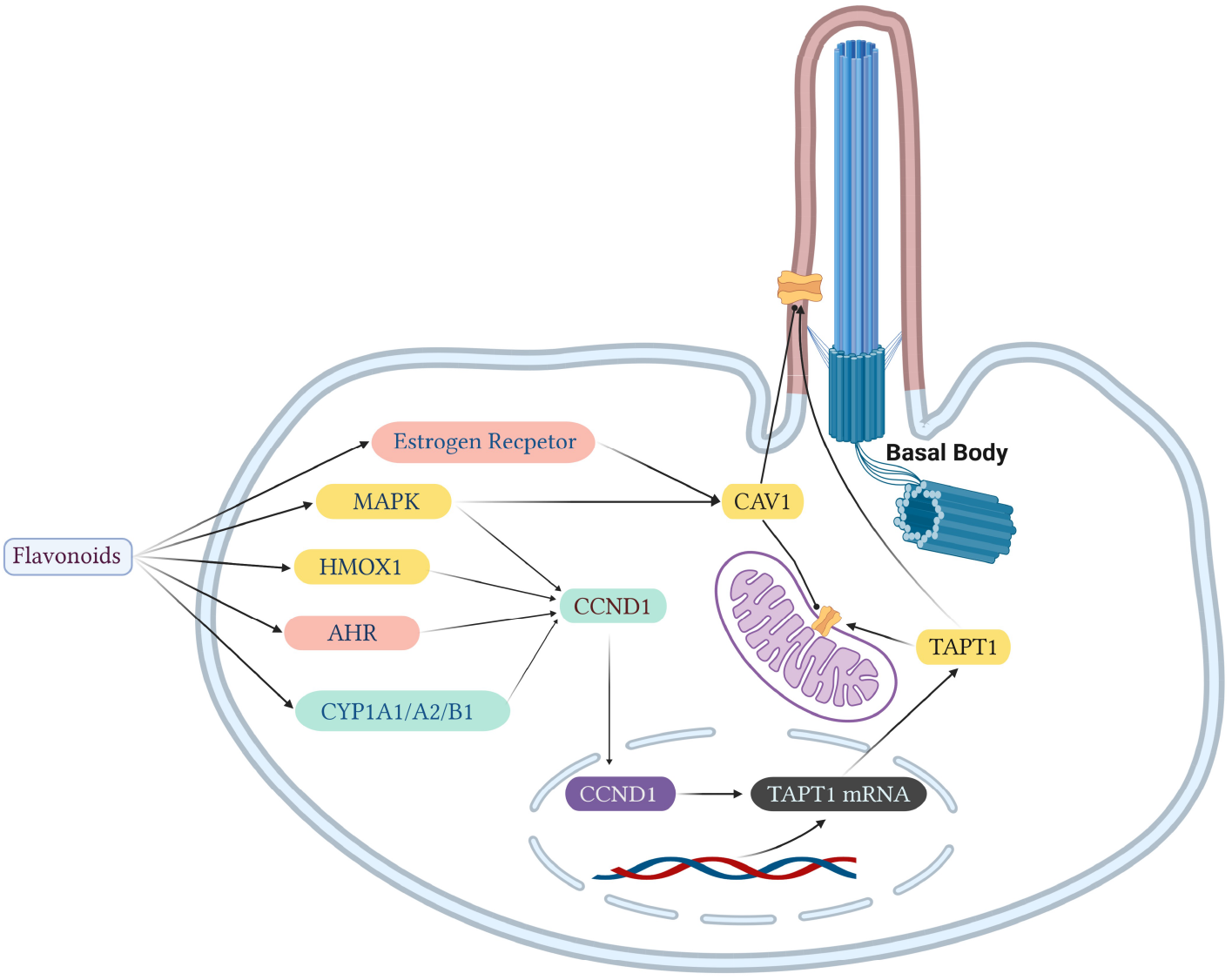
Hypothetical pathway of TAPT1. Here, arrow line represents the increasing of the expression while dot-headed line represents binding. Additionally, yellow color stated for membrane proteins, pink for receptor, cyan for proteins in cytosol and purple for nucleoplasm.

## 4. Discussion

TAPT1 associated with several diseases, including Osteochondrodysplasia, Complex Lethal, Symoens-Barnes-Gistelinck Type, and Microcephaly caused by a mutation in this protein [3]. Nevertheless, it is still unclear how the mutation in TAPT1 leads to these diseases. It is essential to characterize TAPT1, unravel its substrate(s), and discovering the involved signaling pathways. However, due to the structural complexities, the experimental studies of the membrane proteins are challenging. Computational methods offer an alternative way of studying these proteins. Several studies have so far indicated the efficiency of computational methods in predicting the accurate structures of membrane proteins [30].

Moreover, it is very time consuming and costly to determine the substrates experimentally due to the complexity of the metabolic pathways. Therefore, computational approaches may help predict the potential substrates, which will lead to a valid experimental hypothesis [31]. In this present study, a 3D model was generated for TAPT1, and computational methods were employed to study the protein structurally and functionally. Molecular docking approaches were used to predict the substrates, and a knowledge-based ingenuity pathway was analyzed.

As no crystallographic structure was available for TAPT1, 3D models were generated, and the best one was selected according to the assessment score of validation tools. The results of the validation tools demonstrated that the selected model was reliable. Topology and secondary structure analysis from the primary sequence revealed that the protein had eight transmembrane helices. These helices were also predicted from the 3D structure of the protein. Moreover, it revealed that TAPT1 was localized in the mitochondrial membrane in addition to the basal ciliary body, which was also supported by protein-protein interaction analysis (PPI). The protein was found to interact with proteins localized in both the mitochondria and plasma membrane along with extracellular regions. Also, this interactomics revealed the involvement of TAPT1 with some other functions, e.g., intracellular trafficking and viral transport, which were already reported in several studies. A study conducted in a mouse model stated that TAPT1 provided the entry site or gH receptor for human cytomegalovirus [3, 6].

Molecular docking analysis revealed that flavonoids glycosides, as well as flavonoids, would be the potential substrates of TAPT1. The binding site analysis revealed that the amino acid residues involved in the interaction with flavonoids were the same as the pore-lining residues. The amino acid residues H235, R323, K443, N446, S447, L450, K453, S454, Y457, K511, N513, D533, K535, D536, and T538 were found to form the for the surface as well as the binding pocket. Moreover, castP server also predicted the same binding pocket in the TAPT1, which provided strong evidence for this hypothesis. Flavonoids are naturally occurring phenolic compounds that are integral components of the human diet. Several studies revealed that they are involved in the enhancement of bone formation and influence the osteogenic differentiation and mineralization. These compounds also play an essential role in neural crest cell differentiation, proliferation, and survival [32–34]. All of which were disrupted by the mutation in TAPT1 revealed by several studies.

Additionally, mutations in TAPT1 leads to several other diseases related to bone development and disruption in the cilia formation [3]. As flavonoids have a role in bone development and osteogenic differentiation, it was hypothesized that mutations in TAPT1 might disrupt the transfer of flavonoids which caused the osteo-related diseases. Moreover, the ingenuity pathway analysis also provided evidence for the relationship between TAPT1 and flavonoids. IPA revealed that flavonoids could be connected with TAPT1 through several pathways.

## 5. Conclusion

Trans-membrane anterior,-posterior transformation protein 1 (TAPT1) is associated with several ciliopathies and osteogenic diseases. This study provides insights into the structural and functional aspects of the protein. It indicates that flavonoids could be the possible substrate(s) of TAPT1. However, extensive wet-lab experiments are required to validate the outcomes of this study.

## Ethical approval

Not required.

## Acknowledgments

We wholeheartedly acknowledge the cooperation of Sirazum Nadia Hoque (Dept. Computer Science & Engineering, International Islamic University, Chittagong) for helping us during the project designing.

## Data availability

All data supporting the findings of this study are available within the article and its supplementary materials

## Author contributions

MH conceived the study. MP and SA designed the study. MP, MR, MM, DR conducted the experiments. MP, SA, MR, MM, DR, and PC analyzed and interpreted the data. MP wrote the original draft of the manuscript. MP, SA, PC, and MH reviewed and edited the final manuscript. All authors approved the final manuscript.

## Supplementary files

Supplementary Fig 1: The binding pocket of TAPT1 predicted by CASTp server

